# Genetics of cell-type-specific post-transcriptional gene regulation during human neurogenesis

**DOI:** 10.1101/2023.08.30.555019

**Authors:** Nil Aygün, Oleh Krupa, Jessica Mory, Brandon Le, Jordan Valone, Dan Liang, Michael I. Love, Jason L. Stein

**Affiliations:** Department of Genetics, University of North Carolina at Chapel Hill, Chapel Hill, NC 27599, USA; UNC Neuroscience Center University of North Carolina at Chapel Hill, Chapel Hill, NC 27599, USA; Department of Biostatistics, University of North Carolina at Chapel Hill, Chapel Hill, NC 27599, USA

**Keywords:** RNA-editing, alternative polyadenylation, quantitative trait loci, genome-wide association studies, missing regulation, neurogenesis

## Abstract

The function of some genetic variants associated with brain-relevant traits has been explained through colocalization with expression quantitative trait loci (eQTL) conducted in bulk post-mortem adult brain tissue. However, many brain-trait associated loci have unknown cellular or molecular function. These genetic variants may exert context-specific function on different molecular phenotypes including post-transcriptional changes. Here, we identified genetic regulation of RNA-editing and alternative polyadenylation (APA), within a cell-type-specific population of human neural progenitors and neurons. More RNA-editing and isoforms utilizing longer polyadenylation sequences were observed in neurons, likely due to higher expression of genes encoding the proteins mediating these post-transcriptional events. We also detected hundreds of cell-type-specific editing quantitative trait loci (edQTLs) and alternative polyadenylation QTLs (apaQTLs). We found colocalizations of a neuron edQTL in *CCDC88A* with educational attainment and a progenitor apaQTL in *EP300* with schizophrenia, suggesting genetically mediated post-transcriptional regulation during brain development lead to differences in brain function.

## Introduction

Genome-wide association studies (GWAS) have detected many genetic loci associated with risk for neuropsychiatric disorders and inter-individual variation in brain structure and other brain related traits^1–6^. The vast majority of brain-relevant trait GWAS loci have been detected within non-coding genomic regions, and do not change protein coding sequence, implying that they may impact traits through gene regulation^7–9^. Expression quantitative trait loci (eQTL) analysis, which statistically tests the effects of genetic variants on gene expression, have been widely studied in adult brain bulk tissue to interpret the function of brain-relevant trait GWAS loci^10–15^. These studies aggregate steady-state expression across all potential isoforms regardless of post-transcriptional expression modulations. While adult post-mortem bulk tissue eQTLs have identified gene regulatory mechanisms for a subset of brain-trait associated loci, many do not colocalize, leaving their gene-regulatory function unknown. This has recently been termed the “missing regulation” problem in the field of functional genomics^16^. One potential solution could be that brain-relevant trait GWAS loci may function as eQTLs detected only under certain conditions including developmental stage, cell-type, or stimuli^16,17^. Consistent with this, eQTL studies performed in fetal brain bulk data^18,19^, and using cell-type-specific approaches during human neurogenesis and adulthood^20^ identified novel genetic loci associated with brain-relevant traits that were not detected in bulk post-mortem adult brain tissue. Although context-specificity has been taken into account in these studies, there are still brain-trait associated loci that are not explained even by context specific eQTLs. While greater power and additional contexts will likely yield additional gene regulatory effects of brain-trait associated loci, an alternative approach is to study genetic variants influencing brain-relevant traits via alteration of other gene regulatory phenotypes beyond overall expression including post-transcriptional regulation^21–23^.

The most commonly studied post-transcriptional modifications, alternative splicing, has been found to be a major contributor to brain-trait variation using both bulk tissue^18,19,24^ and specific cell-types^25,26^. Many other post-transcriptional events exist, including RNA-editing and alternative polyadenylation, and their genetic regulation are in general poorly studied, especially during the process of neurogenesis. RNA-editing is nucleotide changes in RNA sequence relative to those encoded by the DNA sequence. RNA editing can alter protein product encoded by mRNAs^27,28^, location of transcripts^29,30^, splicing^31,32^, transcript stability^33,34^, and microRNA binding sequences^35,36^. In humans, adenosine-to-inosine (A-to-I) changes are the most common type of RNA-editing^37,38^. A-to-I editing events largely overlap with Alu repeats in the human genome and are generated when ADAR enzymes bind to double-stranded RNA hairpins generated by inverted repeat Alus (IRAlus)^22,39,40^. A-to-I RNA-editing occurs in the human brain^41,42^, and has been shown to impact neurotransmission and neurodevelopment^27,28,43–45^.

RNA-editing dysregulation has been also associated with neurological disorders including schizophrenia^46,47^, autism spectrum disorder^48,49^, major depression^50,51^, epilepsy^52^, and Alzheimer’s disease^53^. Genetic regulation of RNA-editing was observed across adult tissues by performing editing quantitative trait loci (edQTL) analysis, where 1.4% of edit sites (within 228 genes) were significantly regulated by at least one genetic variant^35,40,46^. As a potential mechanism for edQTLs, genetic variants can perturb RNA secondary structure, which can lead to differential RNA-editing in nearby regions^22,35,40^ . Though adult brain bulk tissues are most commonly used to study edQTLs^35,46,54^, recent studies have shown that RNA-editing increases from development to adulthood in the human brain^41,55^, and edQTLs that were not observed in adult brain bulk tissue were detected in bulk fetal brain tissue^55^. Despite the discovery of temporal-specific edQTLs, the cell-type specificity of RNA-editing events and their genetic regulation are as yet unknown during human cortical development and may have been masked in previous bulk tissue studies due to heterogeneity in cell-type composition.

Alternative polyadenylation (APA) is another post-transcriptional modification occurring at the 3’UTR or introns of a given gene in which the poly(A) tail of transcribed mRNA is added in different genomic locations^56,57^. This modification can influence a variety of cellular processes including gene expression, RNA stability, localization, translation rate, and inclusion of microRNA target sequences^56,57^. Dysregulated APA has been found in a variety of brain-relevant diseases including Amyotrophic lateral sclerosis^58^, Parkinson’s disease^59^ and Huntington’s disease^60^. Furthermore, the disruption of several APA regulators have been reported in Oculopharyngeal muscular dystrophy^61^ and multiple neuropsychiatric disorders^62^.

Previous research described the dynamics of alternative polyadenylation (APA) sites during neuronal differentiation and detected that longer 3’UTR isoforms are abundant in neurons compared to progenitors^63–65^. Alternative polyadenylation quantitative trait loci (apaQTL also known as aQTL) analysis can be applied to assess the genetic alteration of alternative polyadenylation site usage^66–69^. Mechanistically, genetic alteration of polyadenylation sites, polyadenylation signal motifs, or motifs of RNA binding proteins (RBPs) regulating APA can impact polyadenylation site usage^66^. apaQTLs performed in human adult brain bulk tissue have identified many genetic loci associated with alternative polyadenylation site usage^68,69^. However, the temporal and cell-type-specific regulation of apaQTLs in the human brain has also as yet remained unexplored.

In this study using an *in vitro* cell-type-specific system model of human neurogenesis, we systematically evaluated genetic regulation of RNA-editing and APA events within progenitors and neurons across ∼80 different donors. We discovered that RNA-editing was more prevalent in neurons compared to progenitors, likely due to higher expression of the genes encoding ADAR2 and ADAR3 proteins. Alternative polyadenylation also showed differences across cell types, where longer 3’ UTR isoforms were observed in neurons as compared to progenitors. We found that common genetic variation was associated with both editing and alternative polyadenylation in a cell-type-specific manner. Hundreds of cell-type-specific edQTLs and apaQTLs were identified that were not previously discovered using fetal bulk brain data. We also observed that these post-transcriptional QTLs, as well as sQTLs showed largely independent regulatory mechanisms as compared to eQTLs. Furthermore, they provided additional interpretation of brain-relevant GWAS loci in addition to eQTLs, suggesting that studying post-transcriptional QTLs is required for a comprehensive understanding of genetic regulation of molecular phenotypes impacting brain development.

## Results

### Cell-type-specific RNA-editing during human neocortical differentiation

We utilized an *in vitro* cell-type-specific system recapitulating human neocortical differentiation in which we previously generated cell-type specific expression, splicing, and chromatin accessibility QTLs^26,70^. This QTL dataset includes progenitors (N_donor_ = 84) and their differentiated, labeled, and sorted progeny, neurons (N_donor_ = 74)^26^. We identified RNA-editing events by observing sequence variants in multiple RNA-sequencing reads (supported by at least 10 RNA-sequencing reads in at least 85% of donors per cell-type) using REDItools software^71^ (Figure 1A and see Methods section). These variants have never previously been identified as genetic variations in human populations. We detected 562 and 3,707 RNA-editing sites in progenitors and neurons, respectively (Table S1). We found that these edit sites overlapped with 261 and 825 genes in progenitors and neurons, and 44% and 59% of genes showed multiple RNA-editing events in those cell types, respectively.

**Figure 1.**
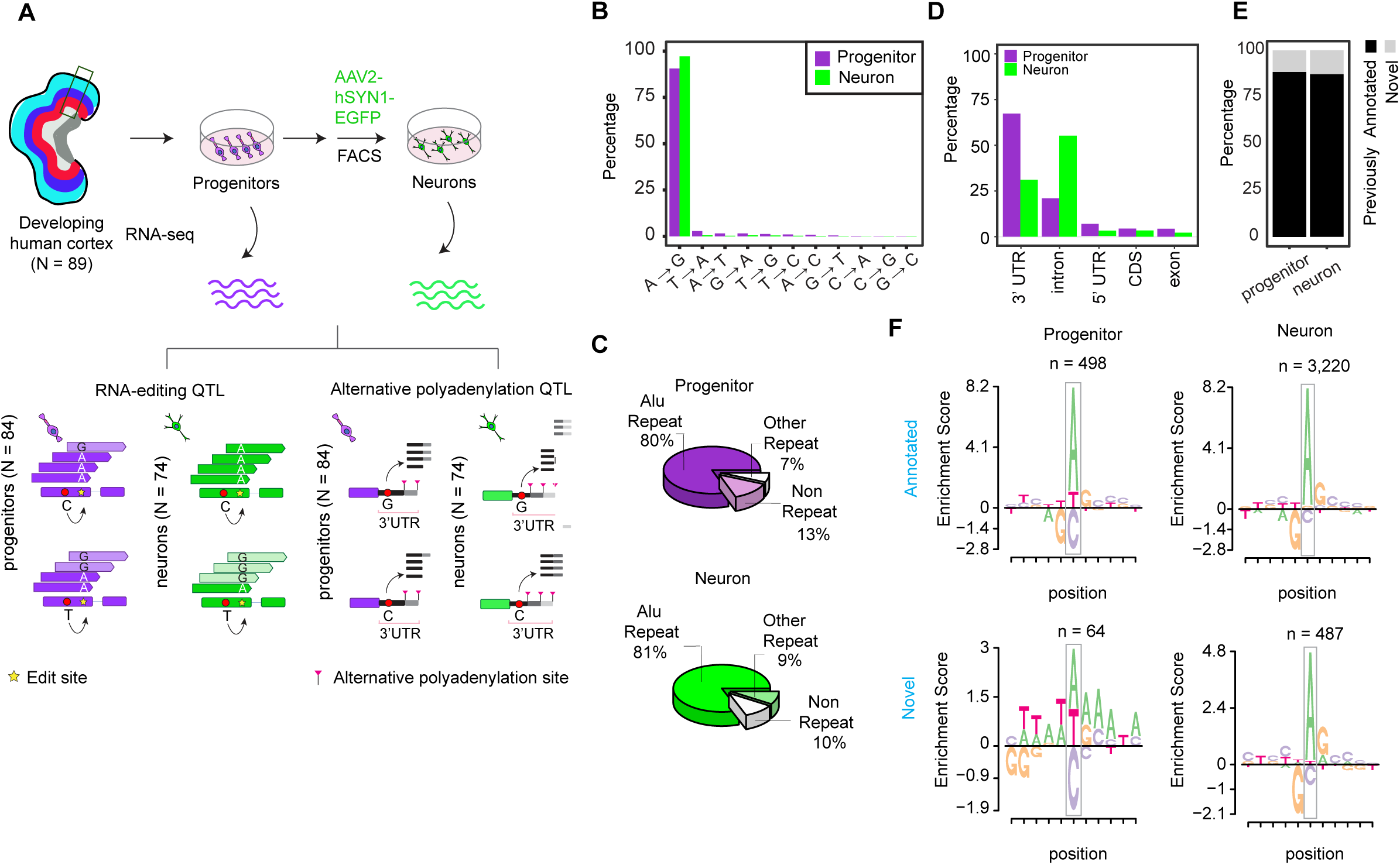
Study design and features of RNA-editing sites. (A) Study design to identify cell-type-specific edQTLs and apaQTLs. (B) Proportions of RNA-editing events discovered in each cell type. (C) Pie chart showing proportions of RNA-editing events within Alu repeats, other repeats or non-repeat regions in the genome per cell type. (D) Proportions of RNA-editing events across genomic regions. CDS: Coding sequence, 3’UTR: Three prime untranslated region and 5’UTR: Five prime untranslated region (E) Overlap of RNA-edit sites with GTEx Cortex or BrainVar datasets. The edit sites overlapped are defined as annotated and the sites which do not overlap are defined as novel. (F) Local motif enrichment for annotated and novel edit sites per cell-type. Number of edit sites (n) in each category is reported.

To validate that these edit sites were not present in genomic DNA, we investigated nucleotides at the genomic position of edit sites within ATAC-seq reads from our previous studies^70,72^. We observed that when any ATAC-seq reads were present at a predicted edit site across all donors, exclusively the unedited allele was present at 94.5% of edit sites, providing confidence that the predicted edit sites were not rare previously undiscovered genetic variants (Figures S1A-B). To evaluate the similarity of RNA-editing sites detected in each cell-type to previously discovered editing events, we examined features including mismatch content and genomic positions of edits. Since the base-pairing features of inosine and guanosine are similar, Adenosine-to-inosine (A-to-I) RNA editing mediated by ADAR is observed as Adenosine-to-guanosine (A-to-G) mismatches in RNA-seq^73^. We observed 90.4% and 97% of mismatches in progenitors and neurons were A-to-G changes (Figure 1B). Most RNA-editing sites in both cell types were overlapped with Alu repeats, also consistent with ADAR mediated RNA-editing (Figure 1C). The majority of RNA-editing was detected within intronic and 3’UTR gene regions, though some was found in the coding sequence (Figure 1D). We also detected that 88% and 87% of RNA-editing sites in progenitor and neurons were also previously identified in either GTEx Cortex^74^ or BrainVar^55^ data in which whole-genome-sequencing data paired to RNA-seq were available (Figure 1E), showing consistency of data generated here with previously discovered RNA-editing events. We further evaluated a common local sequence motif for RNA-editing, which is 1bp upstream enrichment and 1bp downstream depletion of guanosine^75^ among the RNA-edit sites we discovered. We observed that both RNA-editing sites found in GTEx or BrainVar data (previously annotated) and editing sites which were not found in previous datasets (novel) showed highly similar, and expected, motif enrichment in neurons (Figure 1F). However, only previously annotated edit sites showed the expected motif enrichment in progenitors, likely due to the smaller number of edit sites detected in this category (Figure 1F). Supporting that these novel edit sites were not false positives, 97% and 99% of novel edit sites in progenitor and neurons did not show edited alleles in any read from the ATAC-seq data (Figure S1A). These results provided evidence that RNA-edit sites explored in our study exhibited characteristics of previously identified RNA-editing events.

Consistent with the detection of more RNA-edit sites in neurons than progenitors, we found that Alu editing index (AEI), which is calculated as the global measurement of A-to-G changes within Alu elements^76^, was significantly higher in neurons than progenitors (Figure 2A). As a potential mechanism leading to global editing differences between cell types, we compared expression of genes encoding ADAR1, ADAR2, and ADAR3 enzymes. We detected that the expression of *ADAR1* was slightly higher in progenitors than neurons; whereas both *ADAR2* and *ADAR3* were strongly upregulated in neurons (Figure 2B). Consistent with this, increased *ADAR1* expression was positively correlated with global editing levels (AEI) only in progenitors, but increased *ADAR2* and *ADAR3* expression was specifically positively correlated with global editing levels in neurons (Figure S2A). Higher *ADAR2* expression and lower *ADAR3* expression were previously found in neurons compared to oligodendrocytes in the adult brain^54,77^, but have not previously been evaluated in progenitors. Also, *ADAR3* has been previously considered as having an inhibitory role in RNA-editing in the adult brain^54,77,78^, but our developmental and cell-type-specific system revealed a positive correlation between AEI and *ADAR3* expression within neurons, suggesting ADAR3 has a unique role during development leading to increased editing in immature neurons. We also detected a smaller AEI index and number of edit sites discovered in fetal bulk data compared to neurons, and 94% and 96% of edit sites discovered in progenitors and neurons were not identified by using fetal bulk brain data (Figures 2A and C). We observed that average read depth was 17.1M ± 5.8 and 99.8M ± 29.8 in fetal bulk and cell-type-specific RNA-seq data, respectively, so the novel cell-type specific editing events may be driven by either the lower read depth of RNA-seq samples in the fetal bulk data limiting our power to discover RNA-editing events or heterogeneity of cell-types in bulk data. Moreover, consistent with the hypothesis that editing increased throughout the development, we also observed higher AEI values in fetal bulk brain samples^18^ at older gestation weeks as neuronal production increased, consistent with increased editing observed in neurons (Figure S2B). Overall, here, we provide evidence that increased expression of genes encoding ADAR enzymes are likely responsible for cell-type-specific global RNA-editing and the higher number of RNA-editing events in neurons during human neurogenesis.

**Figure 2.**
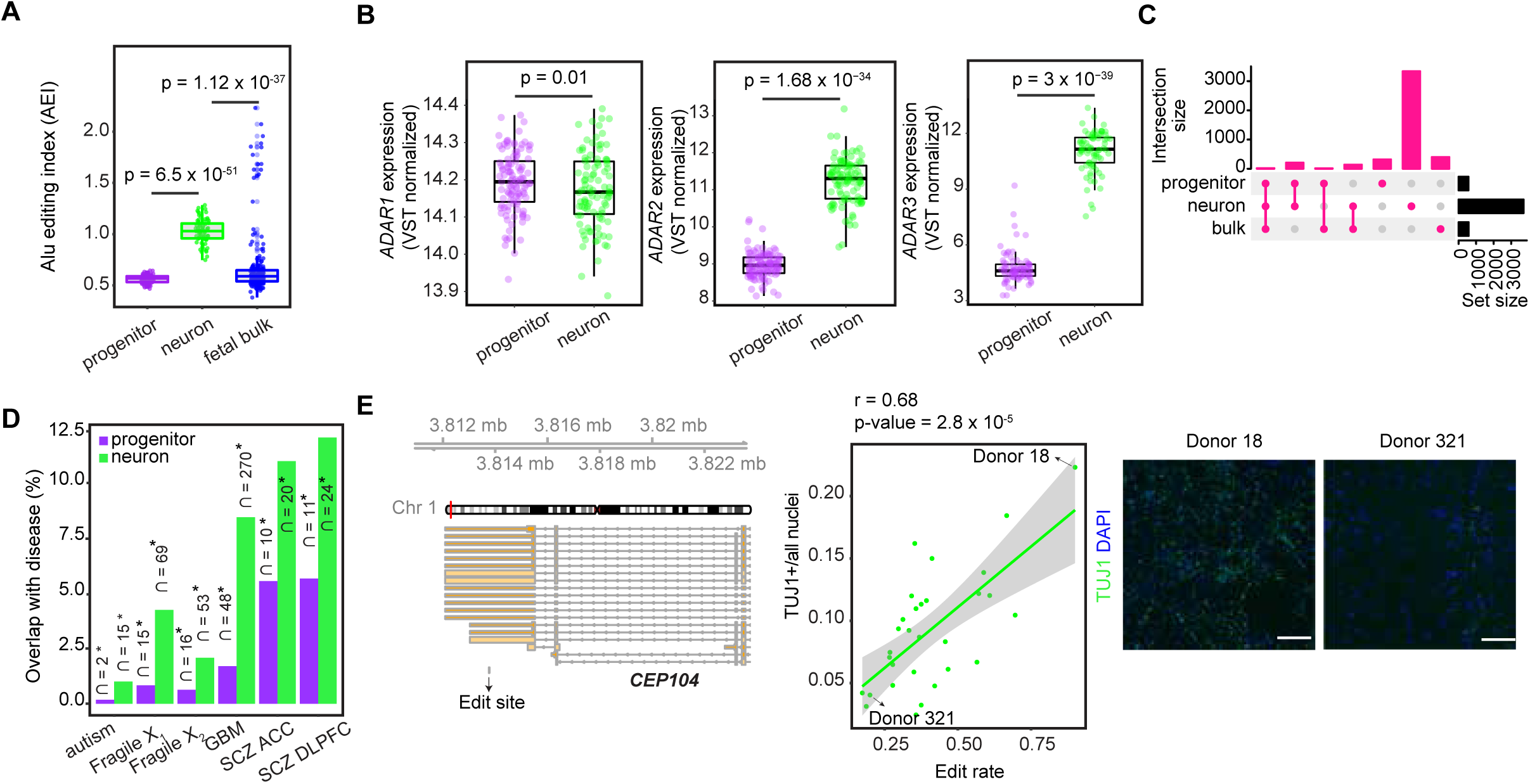
Cell-type-specific RNA-editing during human neurogenesis. (A) Comparison of Alu editing index (AEI) between progenitors and neurons. T-test p-value was reported. (B) Differential expression of *ADAR1*, *ADAR2* and *ADAR3* genes between progenitors and neurons. Adjusted p-value (adj.pval) and log fold change (logFC) from limma were reported. (C) Overlap of RNA-editing sites discovered in progenitors, neurons and fetal bulk data. (D) Enrichment of cell-type-specific RNA-editing sites within edit sites dysregulated in brain-relevant diseases. The number of edit sites overlap is reported, and the proportion of disease-specific edit sites overlapped with cell-type-specific edit sites is shown on the y-axis. Asterisks indicate significant enrichment (FDR < 0.05). Autism (Autism Spectrum Disorder), Fragile X1 (Fragile X syndrome from UC Davis database), Fragile X_2_ (Fragile X syndrome from NIH biobank dataset), GBM (Glioblastoma), SCZ ACC (Schizophrenia from anterior cingulate cortex), and SCZ DLPFC (Schizophrenia from dorsolateral prefrontal cortex). (E) Editing rate of the edit site (chr1:3813767:A>G) at the 3’UTR of *CEP104* gene was positively correlated with the proportion of TUJ1+ neurons. Gene model for *CEP104* and the genomic position of the edit site are given at the left. Scatter plot illustrating the correlation between edit rate and the relative abundance of TUJ1+ neurons, and correlation coefficient (r) and p-values are shown. Representative immunocytochemistry images for TUJ1 (in green) and DAPI (in blue) staining of Donors 18 and 321 (D18 and D321) are shown, scale bar is 100 μm.

Next, we evaluated the downstream functions of RNA-editing. To understand if RNA-editing sites that we discovered were also dysregulated in neuropsychiatric disorders, we evaluated enrichment of cell-type-specific edit sites detected in our sample within previously identified disease-related edit sites which were differentially detected between individuals from case and control groups using adult-bulk post-mortem tissue^46,49,79^. We found that cell-type-specific RNA-edit sites present during neurogenesis that were observed in our model system overlapped with disease-related edit sites found in schizophrenia, glioblastoma, Fragile X, and autism. We observed a greater overlap of disease associated editing events with neurons as compared to progenitors, but these overlaps in both progenitors and neurons occurred more than expected by chance (Fisher’s exact test, FDR < 0.05). These findings suggest a developmental and cellular origin of RNA-editing dysregulation in neuropsychiatric disorders (Figure 2D).

We also evaluated whether RNA-editing influences other cellular downstream functions during neurogenesis. We performed high-content imaging of 8-week differentiated neuronal cultures and labeled them with markers of neuronal differentiation (TUJ1 labeling was used for early born neurons) and all nuclei (DAPI) with 31 wells measured per donor on average. We implemented an image analysis pipeline to quantify the percentage of cells labeled with TUJ1 as a measure of that donor’s neurogenic potential, observing strong differences across donors (compare Donor 18 with Donor 321 in Figure 2E). We found that an editing site within the 3’UTR of *CEP104* gene in neurons was positively correlated with the number of cells expressing a neuronal marker TUJ1 (Figure 2E, Table S2). *CEP104* gene encodes a ciliary protein and loss-of-function mutations on CEP104 were found in individuals with a neurodevelopmental condition, Joubert syndrome^80–82^. These results indicate that RNA-editing can influence fate decisions even without having an effect on the amino acid sequence of the protein.

### Genetic regulation of cell-type-specific editing via editing quantitative trait loci (edQTL) analysis

To perform cell-type-specific edQTL analysis, we tested the association of genetic variants with edit rate, which was defined as the read counts supporting the edited allele (mainly the G nucleoside) divided by the total read coverage at the edit site, within +/-100 kb from edit sites. We included only variants and edit sites located in the same gene (excluding intergenic variants) because we hypothesized that alterations in mRNA secondary structure alter editing. We controlled population structure and global editing principal components (PCs) as technical confounders. We controlled for one global PC of gene editing in progenitors and the major known technical confounder (FACS sorting) in neurons (see Methods), which were highly correlated to *ADAR1-3* expression (Figure S2C). We implemented a hierarchical multiple comparisons correction method using eigenMT-FDR at 5% as a significance threshold (see Methods). We identified 101 and 517 edSites, which are the edit sites significantly regulated by at least one genetic variant, within 79 and 279 genes in progenitors and neurons, respectively (Figure 3A, Table S3). Primary edSNPs, which are variants showing the most significant association with edSites, showed stronger associations as they were closer to the edit sites (Figure 3B). We also observed that genes harboring these edit sites in neurons were significantly enriched in biological pathways related to neuronal morphology and metabolism; whereas we did not detect any significant enriched biological pathways in progenitors (Figure 3C). To investigate a potential mechanism whereby genetic variants impact RNA-editing, we assessed the RNA-secondary structures where significant and nonsignificant edQTLs were located. We found that the majority of significant edQTLs were found within the double stranded RNA-secondary structure, stem, which is substrate for ADAR enzymes in both cell types (Figure 3D). Only significant neuron edQTLs were significantly more enriched within stem structure compared to nonsignificant neuron edQTLs. We did not observe an enrichment in progenitor edQTLs within the stem structure, though this was likely driven by fewer edQTLs discovered in progenitors (Figure 3D).

**Figure 3.**
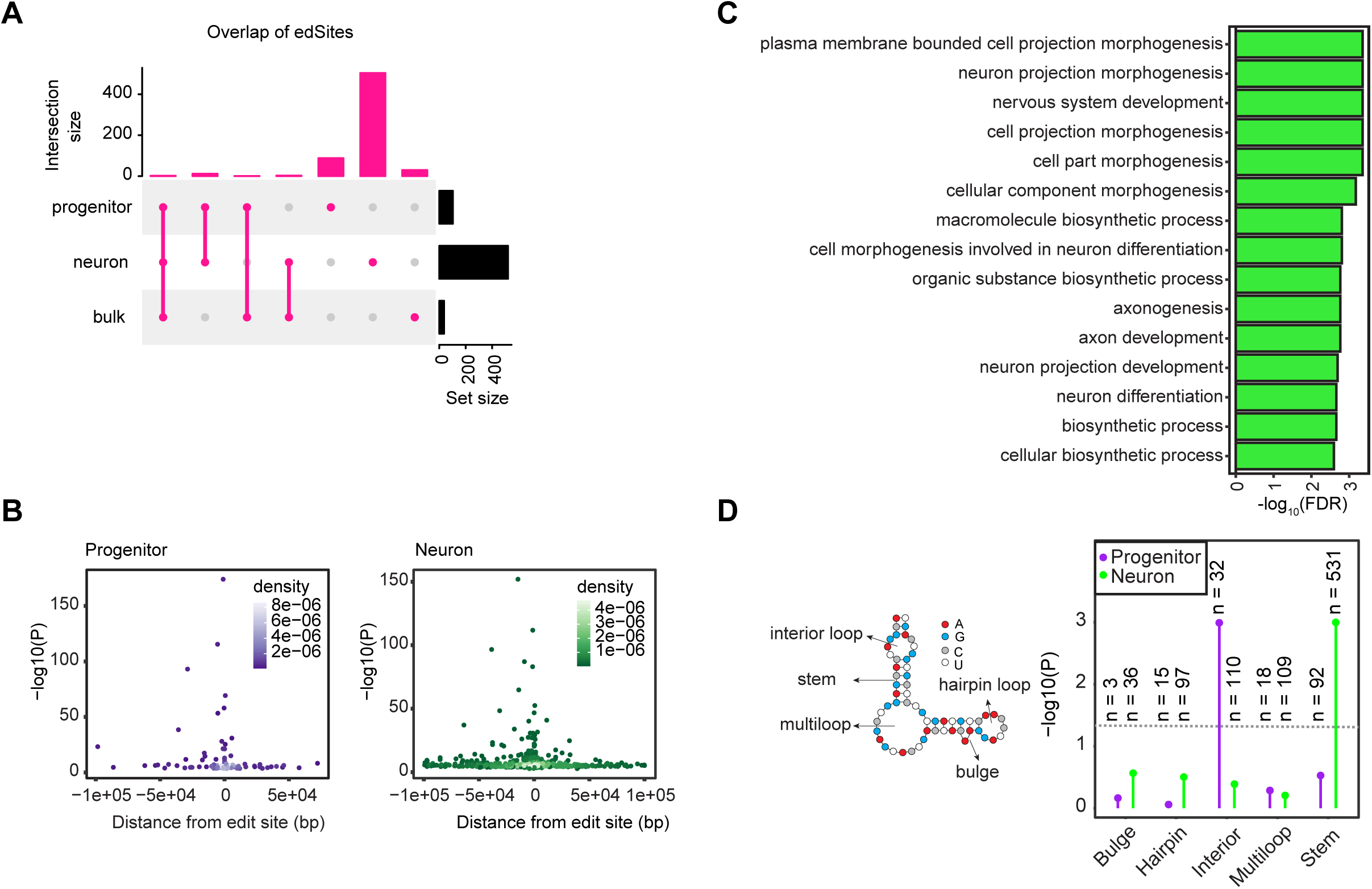
Cell-type-specific edQTLs. (A) Overlap of edSites detected in progenitors, neurons, and fetal bulk data. (B) Primary edQTLs were more significantly associated with editing as they were closer to the edit sites. Association p-values at -log10 scale are shown on the y-axis and distance from edit sites are shown on the x-axis for progenitors (left) and neurons (right), density of the data points are indicated by color density per cell-type. (C) Gene ontology results for edGenes found in neurons. (D) Enrichment of significant edQTLs within different RNA secondary structures illustrated at left. Enrichment p-values at -log10 scale are shown on the y-axis across structures and data are colored by cell-types. The number of variants (n) significantly associated with edit sites and located within each structure are shown.

We observed high cell-type specificity of genetic effects on editing, finding that 86% and 97% of edSites in progenitors and neurons were not detected in other cell types. We also found that 97% and 99% of edSites in progenitor and neurons were not detected in bulk fetal brain tissue, showing additional genetic discovery is enabled using a cell-type specific approach (Figure 3A).

### Cell-type-specific alternative polyadenylation sites during human neocortical differentiation

Another post-transcriptional modification of interest during differentiation is alternative polyadenylation. We identified and quantified alternative polyadenylation (APA) site usage by applying the QAPA method, which allows identification and quantification of 3’UTR isoforms mapped to annotated polyA sites by using RNA-seq data^65,83^ (see Methods). After filtering out lowly expressed 3’UTR isoforms, we detected 19,200 and 18,246 3’UTR isoforms corresponding to alternative polyadenylation sites within 7,711 and 7,801 genes in progenitor and neurons, respectively (Table S4). We observed 2.5 and 2.3 different 3’UTR isoforms per gene on average in progenitors and neurons, respectively. Principal component analysis for APA usage showed that progenitors and neurons were distinctly separated, indicating that cell type has a strong influence on 3’ UTR isoform usage (Figure 4A). We also observed that genes which play a role in alternative polyadenylation^56^ were differentially expressed across cell-types, including higher expression of *FIP1L1* and *RBBP6* in neurons, suggesting their distinct regulation and function during differentiation (Figure 4B). We next performed differential isoform usage analysis across cell-types, and identified both cell-type-specific 3’UTR lengthening and shortening. We observed that the longest 3’UTR isoform of a gene was upregulated in neurons for 79% of 3’UTRs genes, consistent with the previous observation that longer 3’UTRs are expressed during differentiation whereas less differentiated and proliferative cell types generally express shorter 3’ UTRs^65,83–85^. As an example, we found that longer 3’UTR of *CALM1* gene was upregulated in neurons (Figure 4C). CALM1 protein is a calcium ion sensor^86^, and the neuron specific longer 3’UTR expression of the *CALM1* gene was previously shown in mice and its deficiency led to impaired neuronal function^87^. In summary, longer 3’ UTRs were observed in neurons as compared to progenitors, some of which have previously been shown to be functional, and which may be mediated by the increased expression of FIP1L1 and RBBP6 polyadenylation factors.

**Figure 4.**
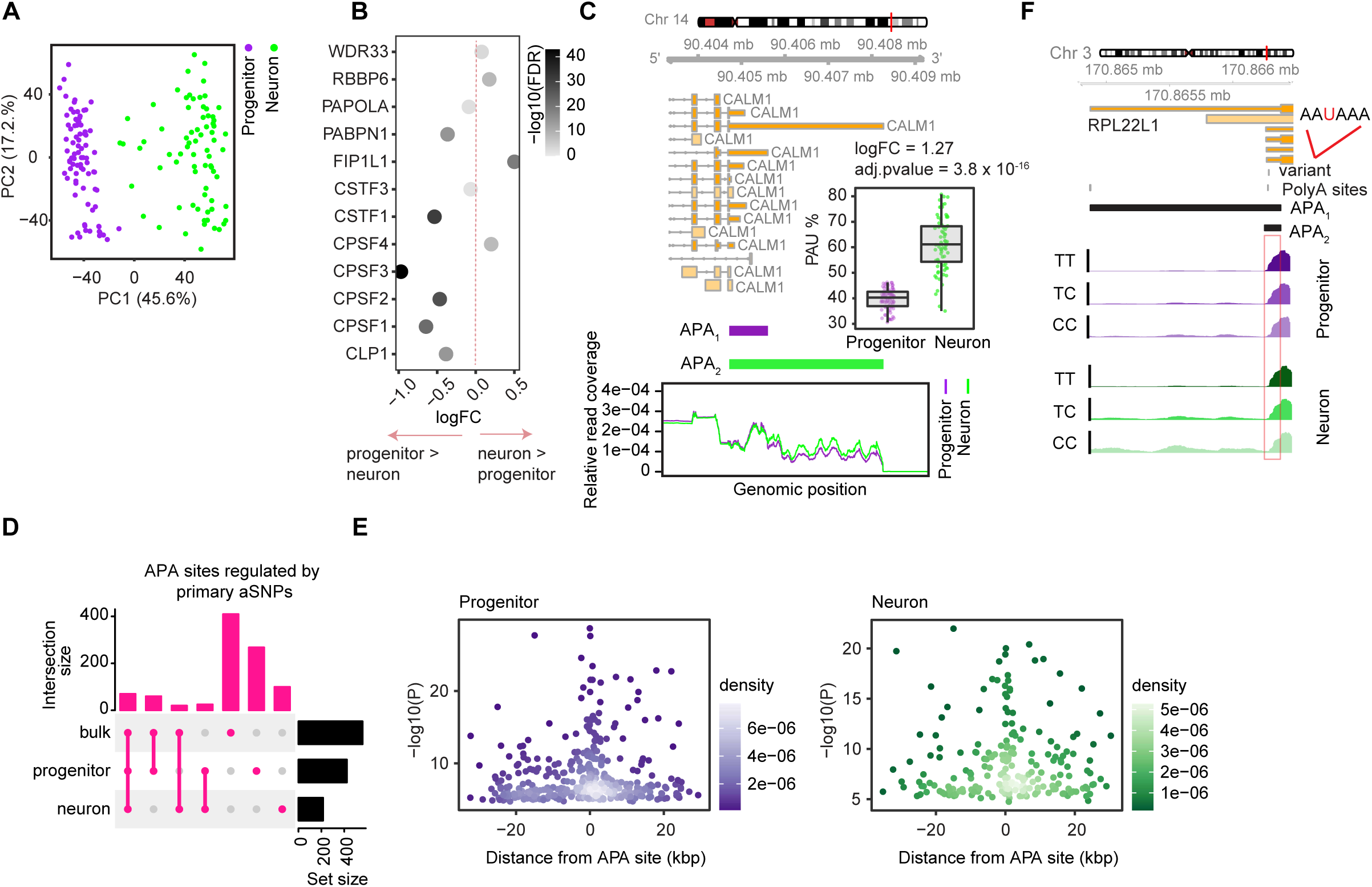
Cell-type-specific alternative polyadenylation and apaQTLs. (A) Principal component analysis for alternative polyadenylation (APA) site usage colored by cell-type. (B) Differentially expressed genes between progenitors and neurons that encode proteins playing a role in alternative polyadenylation. Fold change (logFC) is given on the x-axis and data points are colored by adjusted p-value at -log10 scale. logFC > 0 indicates the genes upregulated in neurons and logFC < 0 indicates the genes upregulated in progenitors, and logFC = 0 is shown with dashed vertical line line. (C) Two APA sites of *CALM1* which were differentially expressed between progenitors and neurons are shown. Gene model for *CALM1* is provided above, and relative read count per APA sites for each cell-type are shown where relative read coverage calculated as ratio of number of count supporting each APA site to total number of reads including both APA sites are shown on the y-axis; and genomic position of the reads are shown on the x-axis. Differential expression of APA_2_ (longer 3’UTR isoform) is shown between cell-types. (D) Overlap of APA sites regulated by primary aSNPs across progenitors, neurons and fetal bulk brain data. (E) Primary apaQTLs were more significantly associated with APA as they were closer to APA sites. Association p-values at -log10 scale are shown on the y-axis and distance from APA sites are shown on the x-axis for progenitors (left) and neurons (right), density of the data points are indicated by color density per cell-type. (F) apaQTL overlapping with canonical polyadenylation signal motif AAUAAA for gene *RPL221* is shown for each cell type. Read coverage per genotype is shown.

### Genetically altered cell-type-specific alternative polyadenylation sites

We performed alternative polyadenylation QTL (apaQTL) analysis by evaluating the association of each 3’UTR isoform with the genetic variants within +/-25 kb window of 3’UTR start and end sites (see Methods). We identified 423 and 215 primary apaQTLs within 352 and 184 genes in progenitors and neurons, respectively (Figure 4D, Table S5). Primary apaQTLs showed stronger associations in closer proximity to the APA site (Figure 4E). To investigate a mechanism underlying apaQTLs, we searched for significant apaQTLs that change canonical polyadenylation signal sequences. We found that 13.6% and 16.3% of significant apaQTLs in progenitor and neurons, respectively, were within canonical polyadenylation signal (PAS) sequences. As previously reported, we also found that AAUAAA was the most frequent motif among these overlapped PAS motifs altered by significant apaQTL^69^. As an example, we detected an apaQTL for *RPL22L1* gene in both progenitor and neuron cells, and the T allele which was the AAUAAA motif matching allele was associated with increase in short 3’UTR and decrease in long 3’UTR (Figure 4F). To evaluate cell-type-specificity of apaQTLs, we utilized π_1_ statistics. We estimated the proportion of progenitor and neuron primary aSNP-APA pairs that were non-null associations (π_1_) in neuron and progenitor apaQTLs as 36.3% and 47.4%, respectively, showing high cell-type specificity of alternative polyadenylation. We next evaluated the overlap of cell-type-specific apaQTLs with fetal brain bulk data, and observed high cell-type specificity of alternative polyadenylation (Figure 4D), again showing that genetic effects on post-transcriptional modifications have greater discoverability within homogeneous cell types.

### Genomic features distinguishing molecular QTLs

Next, we evaluated the genomic features distinguishing different types of molecular QTLs in order to understand their shared or unique regulatory mechanisms. We evaluated this using molecular QTLs previously identified in the same population of neural progenitor cells (expression, splicing, and chromatin accessibility) together with those identified in this manuscript (editing and polyadenylation)^26^. We observed that both primary edQTLs and apaQTLs were more often near transcription termination sites (TTS) as compared to primary eQTLs which were more often near transcription start sites (TSS) in both cell-types (Figures 5A-B). Unlike primary sQTLs, both primary edQTLs and apaQTLs were found less often near splice sites (Figure 5C). Functional genetic variants are often near the molecular entity they regulate.

**Figure 5.**
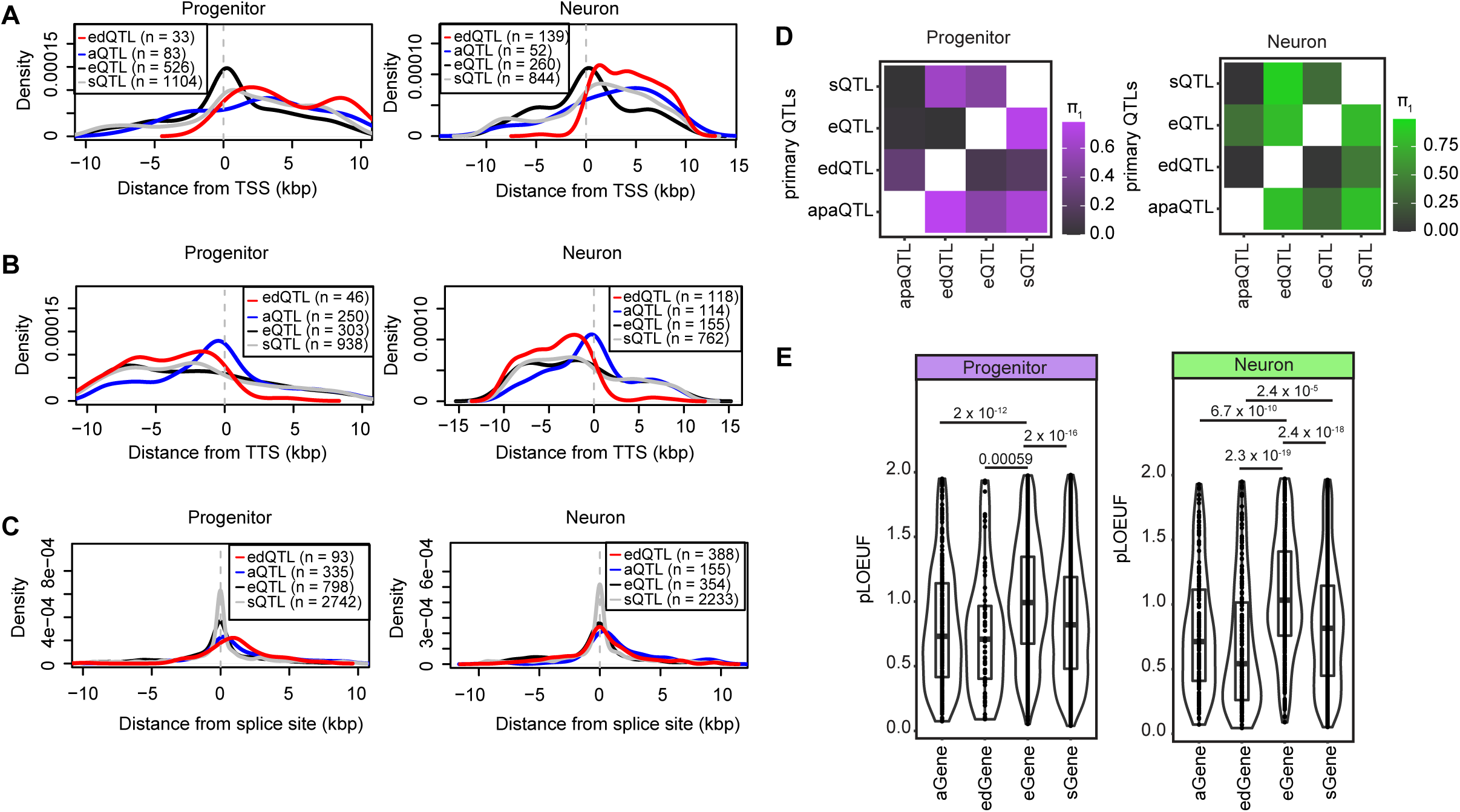
Comparison of cell-type-specific molecular QTLs. (A) Distribution of primary eQTLs, sQTLs, edQTLs and apaQTLs from transcription start site (TSS) per cell-type, the distance from TSS is shown on the x-axis. (B) Distribution of primary eQTLs, sQTLs, edQTLs and apaQTLs from transcription termination site (TTS) per cell-type, the distance from TTS is shown on the x-axis. (C) Distribution of primary eQTLs, sQTLs, edQTLs and apaQTLs from splice sites per cell-type, the distance from splice is shown on the x-axis. Two distances were calculated relative to intron start and end sites, and the shortest distance was used for comparison for each QTL data. (D) Overlap of primary e/s/ed/apaQTLs via π_1_ statistics (progenitors in purple and neurons in green). Matrices are colored based on the proportion of progenitor and neuron primary edSNP-edGene, aSNP-aGene, sSNP-sGene, eSNP-eGene pairs that were non-null associations (π_1_) in each of QTL datasets. (E) pLOUEF values for aGenes, edGenes, eGenes and sGenes per cell type are shown. P-values from t-test are reported.

Furthermore, we also detected that primary edQTLs were found less often within chromatin accessible regions more accessible within that cell type as compared to eQTLs^70^ (Figure S3A, left side). Also, primary apaQTLs were found less often within chromatin accessible regions compared to eQTLs in progenitor cells (Figure S3A, left side). On the other hand, both primary edQTLs and apaQTL were enriched more within RNA Binding Protein (RBP) binding sites^88^ than eQTLs in both cell types (Figure S3A, right side). These findings are consistent with the predicted mechanism of action of eQTLs, alterations in regulatory element activity marked by accessible chromatin, as compared to ed/apaQTLs which are likely dependent on alterations in affinity of RNA binding proteins.

Also, we observed that 2% and 1.8% of edSNPs, variants significantly associated with at least one edit site, were also significantly associated with the expression of the gene harboring the edit sites (edGene) in progenitors and neurons, respectively. 5.8% and 5.6% of apaSNPs significantly associated with at least one APA, were also significantly associated with the expression of the gene harboring the 3’UTR (aGene) in progenitors and neurons, respectively. Given the difference in statistical power between different datasets, we also applied π_1_ statistics^89^ to assess the overlap across edQTLs/apaQTLs/eQTLs and sQTLs. We found that the proportion of progenitor and neuron primary edSNP-edGene pairs that were non-null associations (π_1_) in progenitor/neuron eQTLs were 12% and 2%; the proportion of progenitor and neuron primary apaSNP-aGene pairs that were non-null associations (π_1_) in progenitor and neuron eQTLs were 50% and 34.6%, and the proportion of progenitor and neuron primary sSNP-sGene pairs that were non-null associations (π_1_) in progenitor and neuron eQTLs were 46.5% and 40.6% (Figure 5D). Taken together, these findings suggested that genetic regulation of RNA-editing, alternative polyadenylation, and alternative splicing site usage were mainly independent from eQTLs, consistent with the observations previously reported^22,67^.

To understand whether the small subset of eQTLs shared with ed/apaQTLs were causally related, we performed mediation analysis^90,91^ for variant-edit-gene and variant-APA site-gene triplets. While we did not detect any variant-edit-gene triplets supporting the causal forward model; we detected 335 and 49 variant-APA site-gene triplets. As an example, we found that an apaQTL in progenitors mediated expression of *CEP250* gene (Figure S3B).

We also compared the pLOUEF scores^92^ of the genes harboring edSites (edGenes) and APA sites (aGenes) in this study and eGenes and genes for which splice sites (sGenes) were found to be significantly regulated in our previous study^26^. We observed that sGenes, edGenes and aGenes showed lower pLOUEF scores than eGenes in both cell types. Lower pLOUEF scores indicate genes that are generally protected from rare damaging variation, suggesting that they are important for diseases. These findings indicate that the genes affected by editing and APA are likely to be more disease relevant (Figure 5E).

### Interpretation of the function of the brain-relevant GWASs using post-transcriptional QTLs

To explain the function of genetic variants associated with brain-relevant traits, we next leveraged cell-type-specific edQTLs and apaQTLs with brain-related trait GWAS. Applying colocalization analysis to 2,260 brain-trait GWAS including neuropsychiatric disorders, brain structure and function, and cognitive performance (see Methods), we identified 6 and 6 GWAS loci-traits pairs colocalized with progenitor and neuron edQTLs, respectively; also we found 6 and 3 GWAS loci-trait pairs colocalized with progenitor and neurons apaQTLs, respectively.

Importantly, we did not detect some of these loci in cell-type-specific eQTL and sQTL analysis^26^, suggesting that our approach to integrate post-transcriptional gene regulatory phenotypes revealed the regulatory mechanism of additional brain-relevant trait GWAS loci (Figure S4A, Table S6). As a specific example for edQTLs colocalized with brain-relevant traits, we observed that a neuron-specific edQTL, rs56320407 was co-localized with an educational attainment GWAS-associated locus (index variant rs2589091, p-value = 3.3 x 10^-8^, LD r^2^ = 0.85 based on European population) within the *CCDC88A* gene locus^93^ (Figure 6A). Importantly, the edQTL was not associated with any significant changes in *CCDC88A* expression showing that edQTL enabled the detection of new brain-related genetic variation (Figure 6A). The edit site (chr2:55406089:A>G, Figure S1B) was within the protein coding sequence of one the isoform of *CCDC88A,* though did not change its amino acid sequence, but overlapped the intronic region of the rest of the isoforms. The T allele of rs56320407 was associated with an increase in RNA-editing and decreased educational attainment (Figure 6B). Both the edit site and the index edQTL variant rs56320407, which is 20 bp away from the edit site, were within an Alu repeat (Figure 6C). We predicted the secondary structure of a potential IRAlu hairpin using an *in silico* analysis separately for T and C alleles of variant rs56320407 using RNA sequence between this Alu site and the closest Alu repeat in the opposite direction^94^ (Figure 6C). The predicted secondary structure of the IRAlu hairpin including T (U in RNA sequence) allele matched with the A nucleotide; whereas structure including the C allele caused a C-A mismatch (Figure 6C). Given that the T allele was associated with higher editing, this observation suggests that the T allele caused formation of an RNA secondary structure substrate which was preferred by the ADAR enzymes that consequently led to higher editing level. CCDC88A is an actin binding protein, and it played a role in axonal development and newborn neuron migration during mouse adult neurogenesis^95,96^. Though the edit site did not lead to amino acid change in the protein or differences in mRNA expression, we suggest that the edit site within the *CCDC88A* gene may impact higher cognitive function via altering mRNA stability and eventually translation of protein during cortex development.

**Figure 6.**
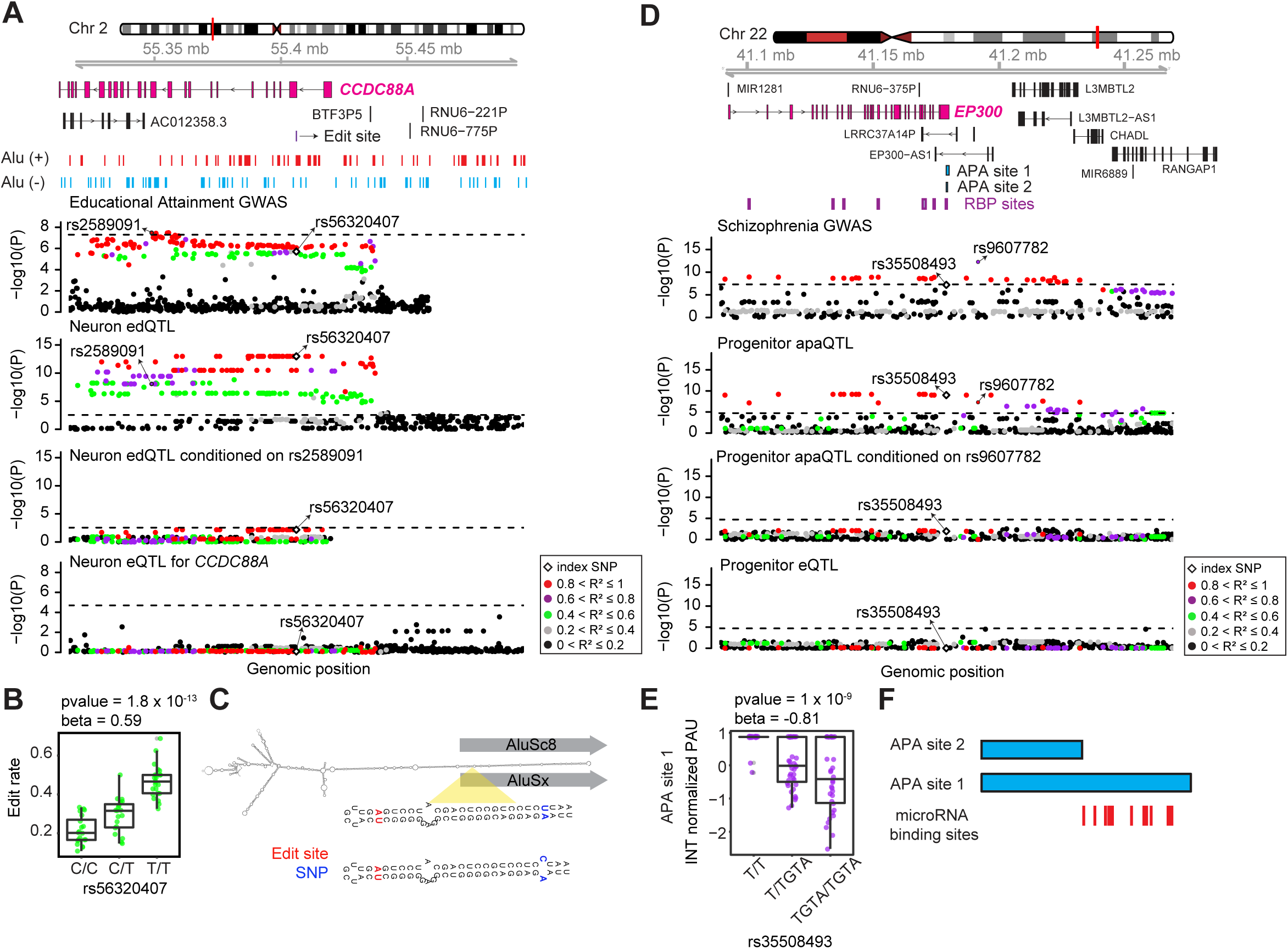
Colocalization of cell-type-specific edQTLs and apaQTLs with brain-relevant trait GWAS loci. (A) Genomics tracks illustrating that an edQTL within *CCDC88A* gene locus in neurons was colocalized with an index variant for education attainment GWAS; whereas there was not any significant eQTL for *CCDC88A*. Data points were colored based on the pairwise LD r^2^ with rs563320407; p-values for each association are shown on the y-axis and genomic positions of genetic variants are shown on the x-axis. (B) Boxplot illustrating distribution of editing rate across rs56320407. (C) The predicted secondary structure of the IRAlu hairpin for T and C alleles of rs56320407. Edit sites and genetic variants are indicated by red and blue colors, respectively. (D) Genomics tracks illustrating that an apaQTL within *EP300* gene locus in progenitor was colocalized with an index variant for schizophrenia GWAS; whereas there was not any significant eQTL for *EP300* gene. Data points were colored based on the pairwise LD r^2^ with rs35508493; p-values for each association are shown on the y-axis and genomic positions of genetic variants are shown on the x-axis. (E) Boxplot illustrating distribution of editing rate across rs56320407. (F) microRNA binding sites are shown for the genomic region which differs between two potential 3’UTR isoforms of *EP300* gene (genomic coordinates for APA site 1: chr22:41,178,956-41,180,079 and APA site 2: chr22:41,178,956-41,179,495).

One example of an apaQTL colocalized with brain-relevant trait GWAS was found at the *EP300* gene locus. We detected that progenitor-specific index apaQTL variant, rs35508493, was colocalized with variant rs9607782 which is an index SNP within the schizophrenia GWAS^3^ (p-value = 5.5 x 10^-13^) (Figure 6D). We did not observe any genetic variants associated with summarized gene expression at the locus. Insertion of GTA nucleotides at rs35508493 was associated with decrease in usage of the longer 3’UTR isoform and lower risk for schizophrenia (Figure 6E). Variant rs35508493 was overlapped with binding sites of multiple RBPs including LIN28B, YTHDF1, YTHDF2, YTDC1 and IGF2BP3 based on CLIPdb database (Figure 6D)^88^, suggesting that it may alter APA site usage by interfering with RBP function. *EP300* gene encodes a histone acetyltransferase protein, and its inhibition promoted proliferation of neural progenitors in adult zebrafish^97^. We observed several microRNA binding sites within the genomic location that differ between long and short APA sites (Figure 6F). This observation suggests that different APA site usage may influence mRNA stability by interfering microRNA function which may consequently lead to differences in protein translation that may influence schizophrenia risk by impacting neural proliferation.

We also evaluated how many additional brain-relevant GWAS loci’s function could be explained via our cell-type-specific system in addition to adult brain eQTLs. We found that our cell-type-specific QTL approach allowed discovering the function of 0.6-4.5% of GWAS loci which could not be explained by adult brain eQTLs previously (Figure S4B). Furthermore, we also investigated that 1.3%-37.5% of adult brain eQTLs which could already explain the function of brain-relevant trait GWAS loci were also cell-type-specific QTLs, indicating a developmental origin of these adult eQTLs (Figure S4B).

## Discussion

In this study, we identified the impact of genetic variants on cell-type-specific RNA-editing and alternative polyadenylation. We found that: (1) RNA-editing was more frequently observed in neurons compared to progenitor cells. This increase in RNA editing may be mediated by higher expression of *ADAR2* and *ADAR3* in neurons. (2) Consistent with previous findings of 3’UTR isoforms lengthening during differentiation^63–65^, the majority of longer 3’UTR isoforms of the genes were upregulated in neurons. (3) Both edQTLs and apaQTLs were strongly cell-type-specific. (4) Both edQTLs and apaQTLs were enriched within genomic regions and regulatory elements which were different from eQTLs, suggesting independent genetic regulatory mechanisms. (5) We found that a few edQTLs and apaQTLs were colocalized with brain-relevant trait GWAS loci in progenitor and neuron cells, increasing the known gene regulatory mechanisms underlying complex brain traits.

Previous studies have reported that RNA-editing increases from fetal to adult human brain^41,55^. However, the mechanism causing the developmental increase of RNA-editing has remained unexplored. Our cell-type-specific design using approximately 80 different donors provided sufficient power to observe the difference in RNA-editing between cell types of the prenatal brain. RNA-editing was six times more frequently observed in neurons compared to progenitor cells during cortical development. This observation was consistent with *ADAR2* and *ADAR3* upregulation, but slight *ADAR1* downregulation in neurons. We observed that higher *ADAR1* expression was associated with increased editing levels in progenitors, but not in neurons. Also, we noted that higher *ADAR2* and *ADAR3* expression were associated with higher editing levels, specifically in neurons. Though increased *ADAR2* was also previously found to be associated with increased editing in the adult brain, *ADAR3* has been previously thought to inhibit RNA-editing using expression measured in the adult brain^54,77,78,98^ . However, here we show that *ADAR3* is positively correlated with editing in neurons suggesting that ADAR3 can act as an activator of RNA-editing specifically during early development, but it may later decrease editing levels in adulthood. Future studies measuring RNA-editing levels following modulation of *ADAR3* in immature and mature neurons will provide insights into addressing this controversy. Increased editing levels in neurons suggests that RNA-editing has functionality in neurons including neuronal differentiation, maturation and activity. For instance, we found that higher editing rate within the *CEP104* gene was positively correlated with higher number of neurons generated during differentiation, while the same edit site was not discovered in progenitor cells (Figure 1E). Also, these results imply that the molecular engineering tools such as RADAR and cellREADR^99,100^ may be more useful in cell types with higher ADAR expression, such as neurons, and that design of sense-edit-switch sequences that target endogenous mRNAs may benefit from knowing which variants increase or decrease editing events.

A previous report indicated that longer 3’UTR isoforms are abundant in neurons compared to progenitors^65^ that 3’UTRs upregulated in neurons were mainly the longest possible isoform of the genes. These observations suggest that testing 3’UTR isoform levels derived from long-read sequencing across different genotypes will also be a useful strategy to validate apaQTLs given that our short-read RNA-sequencing data may have limited accuracy for quantification of the reads mapping to multiple isoforms. Finally, we detected that apaQTLs showed a higher overlap with eQTLs compared to the overlap of edQTLs with eQTLs. Different microRNA binding sites may exist more likely across different APAs which may eventually impact regulation of gene expression via those microRNAs. On the other hand, only single base changes via RNA-editing might not be sufficient to alter microRNA binding sites whereas they can still influence RNA secondary structure and eventually translation of proteins. Performing ribonucleoprotein immunoprecipitation assays^101^ at candidate genetically altered APA and edit sites for the microRNAs will help to explore these potential molecular mechanisms in the future.

Our cell-type-specific approach also increased discovery of genetic regulation on post-transcriptional modulation during human brain development. We observed many more ed/apaQTLs in our cell type specific dataset as compared to fetal bulk brain dataset. This could be due to homogeneous cell populations yielding more accurate quantification of post-transcriptional phenotypes, whereas bulk populations intermingle multiple different cell types each of which has different mechanisms. However, it is also important to note that the lower read depth difference in fetal bulk data compared to the cell-type-specific data might have impacted the number of QTLs discovered. Future studies with greater cellular resolution will likely yield greater discovery of genetically altered post-transcriptional gene regulation.

We observed a very low overlap between edQTLs/apaQTLs and eQTLs, suggesting their impact is independent of the genetic regulation of gene expression as also proposed by several previous reports^22,67–69^. This observation suggests that edQTLs/apaQTLs may impact protein abundance, localization, or function rather than mRNA expression levels. The influence of APA on protein abundance and ribosome occupancy without altering mRNA expression levels has been previously described^67^. Although there is no clear evidence for the impact of RNA-editing on translation yet, changes in protein levels via RNA-editing were detected in a previous study^102^. Previous studies in adult brain data showed the differences in the impact of genetic variants on gene expression and proteins^103–105^. Comparison edQTLs/apaQTLs with protein QTLs in cell-types of developing brain will clarify what the functional consequences of these variants are at the molecular level.

As a result of their independent regulation, novel post-transcriptional QTL colocalizations can be detected with brain-relevant trait GWAS that were not observed in steady state eQTL colocalizations. Importantly, we observed a few edQTLs and apaQTLs colocalized with brain-relevant trait GWAS, contributing to solving the missing regulation problem. Utilization of edQTLs/apaQTLs with larger sample size may enable explanation of additional genetic mechanisms underlying complex brain traits in the future.

## Acknowledgements

This work was supported by NIH (R00MH102357, U54EB020403, R01MH118349, R01MH120125). The following core facilities were utilized for this project: UNC Neuroscience Center Microscopy Core (P30NS045892), UNC Mammalian Genotyping Core, CGIBD Advanced Analytics Core (NIH grant P30 DK034987), UNC Flow Cytometry Core Facility, UNC Vector Core, UNC Research Computing. Additional core facilities utilized for this project were: UCLA CFAR (5P30 AI028697), and the UCLA Neuroscience Genomics Core.

## Author Contributions

JLS and NA conceived the study. JLS directed and supervised the study. JLS provided funding. DL, BL and JV processed ATAC-seq data. OK and JM performed immunocytochemistry. JM and NA quantified marker expression. MIL aided in editing quantitative loci methodology. NA managed the integration of the datasets, interpreted results, performed genotype imputation, identified RNA-editing and alternative polyadenylation sides, performed edQTL, aQTL, co-localization, mediation, and enrichment analyses. JLS and NA wrote the manuscript. All authors commented on and approved the final version of the manuscript.

## Availability of data and materials

Codes are available here https://bitbucket.org/steinlabunc/post_transcriptional_qtls/src/master/

**Figure S1. Validation of RNA-editing sites discovered in RNA-seq data via ATAC-seq data.**

(A) The number of donors supporting that genomic DNA includes unedited alleles are shown on the x-axis. The percentage of unedited allele per edit sites is shown on the right y-axis (red) and the total number of ATAC-seq supporting unedited allele is given on the left y-axis (blue).

(B) Example ATAC-seq reads for RNA-editing sites chr19:18365401:A>G, chr6:163422494:A>G and chr2:55406089:A>G.

**Figure S2. ALU editing index is correlated with ADAR gene expression and developmental time.**

(A) Correlation of AEI values with VST normalized ADAR1, ADAR2 and ADAR3 expressions in progenitors (purple) and neurons (green). Correlation coefficients (r) and p-values are shown per relationship.

(B) Alu editing index across different gestational weeks in fetal bulk brain data.

(C) Correlation of technical confounders with *ADAR1-3* gene expressions per cell-type.

Correlation efficient (r) and p-values are reported.

**Figure S3. Comparison of cell-type-specific molecular QTLs**

(A) Overlap of primary eQTLs, sQTLs, edQTLs and apaQTLs with chromatin accessibility regions are RBP binding sites per cell type. Chi-square test p-values (p) are reported for each pairwise comparison.

(B) APA site (chr20:35511626-35519280) within the *CEP250* gene mediated its expression. Genomic tracks for progenitor eQTLs and apaQTL are shown, and tracks are colored based on relative LD r^2^ for the variant rs2236160, which is the nearest variant to APA site.

(C) Posterior probability of forward, independence and reactive models for mediation of *CEP250* gene expression via APA site. Lower cartoon illustrates the relationship between variant (X) within the APA site, mediator APA site (M) and *CEP250* gene.

**Figure S4. Comparison of GWAS colocalizations across cell-type-specific molecular QTLs**

(A) Number of brain-relevant trait-loci pairs are found in eQTLs, sQTLs, edQTLs and apaQTLs, and shared pairs with eQTLs are indicated by black color, and pairs which were detected by using QTLs other than that eQTLs are indicated by blue color.

(B) Number of brain-relevant trait-loci pairs are found in cell-type-specific eQTLs, sQTLs, edQTLs and apaQTLs in addition to loci explained by adult brain eQTLs.

## Methods

### Preparation of primary human neural progenitor cells (phNPCs)

We established phNPCs culture including neural progenitor cells and neuronal progeny differentiated from these progenitors by following the experimental workflow described in our previous work^26,70,106^. We acquired the human fetal brain tissue (14-21 gestation weeks old) derived from voluntary terminated pregnancy according to the IRB regulations at UCLA through the Gene and Cell Therapy Core. The tissue pieces corresponding to the cortex were visually selected for generation of phNPCs. In the Geschwind lab at UCLA, we dissociated these tissues and formed neurospheres by using them as we have previously described^26,106^. We then plated the neurospheres on the plates coated with laminin/fibronectin and polyornithine, and after an average of 2.5 +/-1.8 standard deviation passages, they were cryopreserved to transfer to Stein lab at UNC-Chapel Hill^26^.

phNPCs from 89 unique donors were further randomly grouped into 8-9 donors for 12 rounds, and each round was thawed every 3 weeks. Each round was processed by mostly the same person and on the same day of the week, where changes in these technical variable changes were documented. We cultured progenitor cells in proliferation media as previously described^26,70^ for three weeks, and then prepared RNA-seq libraries. To differentiate progenitor cells into neurons, we first cultured the cells in the media without growth factors for 5 weeks and then transduced cells with AAV2-hSyn1-eGFP virus (with 20,000 multiplicity of infection (MOI)) that carries a reporter gene expressed specifically in neurons^26^. After viral transduction, we differentiated the cells for 3 weeks longer, and isolated EGFP-labeled neurons using FACS sorting machines BD FACS Aria II or Sony SH800S. We kept the neurons within the Qiazol solution to prepare RNA-seq libraries.

### RNA-sequencing and data processing

We prepared RNA-seq libraries and sequenced them as described in our previous work^26^. We obtained 150 bp paired end reads with a mean read depth of 99.8M ± 29.8 SD read pairs per library.

To process the RNA-sequencing data, as described in our previous work^26^, if the same library was sequenced on multiple flow cells, we merged .fastq files, trimmed the adapters for all libraries using Cutadapt/1.15 software^107^, and performed quality control with FastQC software.

We aligned the RNA-seq reads into GRCh38 release92 reference genome including sequence of AAV2-hSyn1-eGFP plasmid using STAR/2.6.0a aligner program^108^.

To assess consistency of genotypes detected by genotyping array and RNA-seq, we performed VerifyBamID analysis (v.1.1.3)^109^, as described previously^26^. We retained the RNA-seq libraries with [CHIPMIX] < 0.04 and [FREEMIX] < 0.04, and assigned correct donor IDs for 8 libraries where there was a sample swap. Additionally, one library was missing cDNA concentration and removed, also libraries with eRIN score lower than 7 were not included.

Following quality control, we obtained 84 and 74 unique donors for progenitors and neurons, respectively. We applied the same workflow to process RNA-seq data derived from fetal bulk brain samples and retained 235 unique donors as previously described^26^.

### Genotyping and imputation

We performed genotyping by utilizing HumanOmni2.5Exome or Illumina HumanOmni2.5 platforms followed by filtering via PLINK v.1.90b3 software^110^ with the parameters –hwe 1 x 10^-6^ –geno 0.05 –mind 0.01 –maf 0.01 as described previously^26,70^. We utilized the TOPMed imputation server for imputation with the TOPMed reference panel (Version R2 on GRC38)^111^, after processing the genotype data via imputation preparation pipeline by using the algorithm *perl HRC-1000G-check-bim_v3.pl -b <BIM file> -f <FREQUENCY file> -r 1000GP_Phase3_combined.legend.gz -g -p ALL* (https://www.well.ox.ac.uk/~wrayner/tools/).

For downstream analyses, we retained the variants with following criteria: minor allele frequency (MAF) > 0.01, imputation quality score R^2^ > 0.3 and Hardy-Weinberg equilibrium at p > 1 x 10^-6^ .

### Detection and quantification of RNA-editing events by using RNA-sequencing data

To detect RNA-editing events by using RNA-sequencing data, we first aimed to reduce mapping bias using the WASP method^112^ and remapped RNA-seq reads mapped via STAR/2.6.0a aligner^108^. Following remapping, we discarded duplicated reads and extracted uniquely mapped reads as input for Reditools software v2.0^71^. Using Reditools software, we identified and quantified RNA-editing sites by applying the following parameters: -S -s 2 -q 20 - bp 25 -ss 5 -mrl 50 -C -T 2 –os 5 where -S was used for including only edit sites in the output, - s was for strand inference, -q was for minimum read quality score, -bp was for minimum base quality score, -ss was for splicing span, -mrl was for minimum read length, -C was for strand correction, -T was for strand confidence and –os was for omopolymeric-span. We used Homo sapiens gene ensembl v.92 as the reference genome. We discarded multi-allelic sites and the sites overlapped with genomic variants from our genotype data with imputation R^2^ greater than 0.3 and common SNPs in dbSNP (v153) unless they were listed in the REDIportal database^74^.

For downstream analysis, we retained RNA-editing sites supported by at least 85% of donors with 10 RNA-seq counts (at least 2 counts for edited allele) per cell-type and fetal bulk tissue. We defined RNA-editing rate as the ratio of the number of read counts supporting the edited allele to the sum of the number of the read counts supporting both edited and unedited alleles.

### Validation of RNA-edit sites via matched ATAC-seq data

To confirm that RNA-edit sites are nucleotide changes in RNA-sequence but not genomic mutations, we extracted WASP-mapped ATAC-seq reads from our two previous studies^70,72^ which overlapped with per edit site. We calculated the abundance of each unedited allele of each edit site per donor by getting the ratio of number of reads supporting the unedited allele to total coverage at that site (Figure S1A). We excluded the nucleotides with base quality lower than 30, and the reads where any mismatch were reported in CIGAR string since they obscure the finding of the nucleotides at the genomic region of interest.

### Calculation of the ALU editing index

We computed Alu editing index (AEI) per RNA-sequencing sample bam file per cell-type and fetal bulk tissue (including uniquely mapped reads after STAR alignment and WASP algorithms as used for the detection of RNA-editing via Reditools) via the RNAEditingIndexer algorithm^76^.

### Enrichment of local motif for RNA-edit sites

For each edit site, we extracted RNA-sequence within +/-4 bp window of the edit sites. Providing these sequences to the EDLogo software, we quantified and visualized local sequence motifs^113^.

### Enrichment of progenitor and neuron RNA-edit sites within disease-relevant edit sites

To assess the significance of overlap between disease-relevant edit sites (defined based on adjusted p-value < 0.05 in differential editing analysis between case and control) and edit sites detected in each cell type, we applied GeneOverlap R package.

### Detection of alternative polyadenylation sites from RNA-seq data for each cell type

To detect and quantify alternative polyadenylation by using RNA-seq data, we followed the pipeline described by QAPA method^65^. We initially built 3’UTR libraries including potential alternative polyadenylation sites for each gene by using biomart ensembl gene metadata table (human version 92), GTF file Homo_sapiens.GRCh38.92 as gene prediction table, and PolyASite database (hg38) and GENCODE poly(A) sites track (hg38) as poly(A) site annotations followed by extraction of 3’UTR sequences by integrating these annotations with reference genome fasta file (GRCh38 release92). After trimming the sequencing adaptors from RNA-seq libraries via Cutadapt/4.1^107^, we quantified 3’UTR isoforms via salmon/1.9.0^114^ based on the 3’UTR library generated by QAPA by correcting for GC and sequence-specific biases. To infer poly(A) usage (PAU) value for a given 3’UTR isoform of a gene, we divided the expression of this 3’UTR isoform by the sum of the expression of all other 3’UTR isoforms detected for that gene via qapa *quant* function^65^. For the downstream analyses, we applied the following steps: (1) we retained the 3’UTR isoforms supported by 10 counts at least 10% of the donors for each cell-type. (2) For the genes in which we detected only two 3’UTR isoforms, we randomly selected one of the isoforms to prevent statistical bias since these two isoforms were complementary to each other. (3) We normalized the PAU ratios for the remaining 3’UTR isoforms via quantile normalization.

### Differential alternative 3’UTR expression analysis

Prior to the differential alternative 3’UTR expression analysis, we first corrected quantile normalized neuron gene expression data for the batch effect caused by the usage of different machines for FACS sorting via *removeBatchEffect* function of limma R package^115^. Followed by batch correction for neurons, we combined two cell-type-specific data, and performed a paired differential gene expression via limma package by using the design matrix model.matrix(∼Cell-type + as.factor(DonorID) + RIN, dataset). We identified differentially expressed 3’UTR isoforms between cell-types as adjusted p-values < 0.05 after multiple test correction via Benjamini-Hochberg method^116^. We assessed the isoform lengthening or shortening based on if the differentially expressed 3’UTR had the longest or the shortest 3’UTR isoform among all potential 3’UTR isoforms for a gene.

### Immunohistochemistry of neurons for TUJ1 marker and quantification

We used the same experimental procedure to generate neurons from phNPCs as described previously^26,70^. At 8 weeks of differentiation, we fixed neuron cells in 4% PFA, and permeabilized them by using 0.4% Triton in PBST solution and performed blocking within 10% goat serum dissolved in PBST. After we incubated primary antibodies for TUJ1 (1:2000, Catalog # 801202) overnight in 3% goat serum dissolved in PBST solution at 4°C, we washed the cells three times with PBST. We applied fluorophore-conjugated secondary antibodies (Alexa Fluor 488, goat anti-mouse, 1:1000, Invitrogen, Catalog # A11001) at room temperature for an hour, and applied DAPI staining for 10 minutes.

We performed imaging by using Nikon Eclipse Ti2 with pco.edge 4.2Q High QE sCMOS camera via 10x objective. Prior to segmentation, we used ImageJ to isolate the DAPI channel, transformed them to grayscale and divided images into 4 crops at 2862 by 2862 pixels. We applied Cellpose software ^117^ for segmentation by implementing a nuclear segmentation method in which we set the nucleus diameter as 9 microns. We subtracted the cell outlines from the generated cell masks resulting in a final nuclear mask. To count TUJ1+ cells, we applied CellProfiler to masks generated by Cellpose. We excluded cells if they had nuclei smaller than 6 microns in diameter, which were likely dead cells. In the other image channels, objects with high intensity were considered debris and masked out of the images to aid in threshold and background intensity calculations. For each channel, we corrected images for illumination inhomogeneity, measured background intensity, and images intensities. We classified each cell for the TUJ1 marker if the average intensities were at least 1.5 standard deviations above the median of the background intensity.

### Cell-type-specific editing quantitative loci analysis

We tested association of editing rate with genetic variants within +/-100 kb window of editing site and located in the same gene harboring edit site to perform editing QTL (edQTL) analysis. We included the editing sites if at least 85% of samples had at least 10 read counts (at least 2 counts to support the edited allele) to support the editing site for each cell type. We retained only the variants if at least two heterozygous donors and at least two homozygous minor allele donors, or no homozygous minor allele donors were present as a filtering strategy we previously used^26^. Since the donors showing the sufficient read counts might be different across editing sites, for each editing site, we used 85% of the donors that every one of them supported editing site with at least 10 counts in each cell type (at least 2 counts for edited allele).

We performed cell-type-specific edQTL mapping analysis by using a generalized linear model with binomial distribution that controls for population stratification and unmeasured technical variation. To control for population stratification, we calculated MDS of global genotype and used the first three MDS components as covariates. We controlled the unmeasured technical variation which affects RNA-editing via an optimization strategy. For each cell-type, we utilized principal component analysis (PCA) for unmeasured technical variation, and computed global editing PCs via prcomp() function from stats R package by using edit rate values per edit site. During the optimization strategy, we re-performed edQTL analysis by sequentially adding the global editing PCs, first 3 MDSs of global genotype for each cell-type. We detected FACS sorter as a major technical factor impacting editing rate in neurons, and controlled for it for neuron edQTL analysis (p-value = 1.8 x 10^-7^ for PC1 of global editing and FACS sorter relationship). After each run, we calculated the number of edit sites significantly associated with at least one genetic variant (edSite) at a 5% false discovery rate. Since we found that including 1 PC of global editing maximized the edSite discovery in both progenitor and neurons:

The optimized model we used for progenitors was:

Edit rate ∼SNP + 3 MDS of global genotype + 1 PC of global editing The optimized model we used for neurons was:

Edit rate ∼SNP + 3 MDS of global genotype + FACS sorter + 1 PC of global editing

As implemented in our previous work^26^, we applied a hierarchical correction procedure termed eigenMT-FDR^118^, which allowed us to stringently control for multiple comparisons by considering both the number of edit sites and the variants tested. In this algorithm, we first computed locally adjusted p-values for cis-SNPs per edit site via the eigenMT approach in which a genotype correlation matrix was used to estimate the effective number of independent tests^119^. Then, we performed FDR procedure by using locally adjusted p-values that resulted in globally adjusted p-values per edit site. As the last step, the edit sites with a globally adjusted p-value lower than 0.05 were defined as edSites. We conducted the same procedure to discover edQTLs in fetal bulk brain data.

### Cell-type-specific alternative polyadenylation quantitative loci analysis

To perform cell-type-specific alternative polyadenylation QTL (apaQTL) analysis, we tested association of quantile normalized PAU values per 3’UTR isoform with the genetic variants within the +/-25 kb window of isoform start and end sites. We retained the genetic variants if they were carried by at least two heterozygous donors and without any homozygous minor allele donors, or if they were carried by at least two minor allele homozygous donors identical to edQTL analysis.

We performed apaQTL mapping analysis by controlling for population stratification and cryptic relatedness via a linear mixed effects regression model by using EMMAX software^120^. We controlled the population stratification by using the first three MDS components of global genotype as covariates. We generated the identity by state (IBS) kinship matrix by implementing emmax-kin -v -h -d algorithm by using variants from the non-imputed genotype data, and excluded the variants on the same chromosome via MLMe method^121^. Similar to edQTL analysis, we performed an optimization strategy to identify the number of PCs of global expression of 3’UTR isoforms, which was computed via prcomp() function of stats R package by using quantile normalized PAU values per edit site separately for each cell type. After sequentially adding PCs of global expression of 3’UTR isoforms as covariates to re-run apaQTL analysis, we identified 9 PCs and 6 PCs of global 3’UTR isoform expression in progenitor and neurons showed the highest number of APA site significant associated with at least one genetic variant at 5% FDR. As a result, we applied the following models per cell-type:

The optimized model we used for progenitors was:

PAU ∼ SNP + 3 MDS of global genotype + 9 PC of global 3’UTR expression + ɛ

The optimized model we used for neurons was:

PAU ∼ SNP + 3 MDS of global genotype + 6 PC of global 3’UTR expression + FACS sorter + ɛ

We conducted same pipeline to discover apaQTLs in fetal bulk data and the optimized model we used for fetal bulk was:

PAU ∼ SNP + 3 MDS of global genotype + 5 PC of global 3’UTR expression + ɛ

where PAU is Poly(A) usage^65^ and we defined an error term ɛ for which cov(ɛ) = σ^2^ *u* + σ^2^ *I* in which kinship matrix used to account genetic relatedness is indicated by *u_K_*, variance is indicated by σ^2^u and σ^2^e is random noise of the variance.

### Assessment of QTL sharing between cell-types and different molecular QTLs

For cell-to-cell comparison, the proportion of progenitor and neuron primary SNP-edit site pairs or APA site pairs that are non-null associations in neuron and progenitor edQTLs or apaQTLs data was estimated by utilizing the corresponding p-values to SNP-edit site pairs or APA via *π*_1_ statistics^89^ by using the qvalue R package^122^. Similarly, for edQTL/apaQTL to eQTL comparison, we estimated the proportion of progenitor and neuron primary SNP-edGene pairs, gene including edit site or SNP-aGene pairs, gene including APA site pairs that are non-null associations in progenitor and neuron eQTL data (*π*_1_) by using the corresponding p-values to SNP-Gene pairs detected in both datasets.

### Prediction of inverted repeat Alus (*IRAlu*) RNA hairpin secondary structures

To predict IRAlu RNA hairpin secondary structure, we extracted RNA-sequences between two Alu repeats in opposite directions, and generated two sequences corresponding to different alleles of a given genetic variant. These sequences were provided to viennaRNA RNAfold software, and secondary structures were predicted and visualized via graphical output from the software^94^.

### Enrichment of edQTLs within RNA secondary structures

As described in previous study^22^, for each edSite (edit site significantly associated with at least one genetic variant), we extracted the RNA sequences including genetic variants within +/-800 bp window of the edSites. We included only sequences within gene start and end coordinates within this window; therefore, some sequences were shorter than 1601 bp. We matched the alleles for variants within this genomic window if the LD r^2^ between them was greater than 0.8. We first converted sequences files to bpseq via contrafold software^123^. Next, providing these bpseq file formats for bpRNA software^124^, we predicted RNA secondary structures including bulge, hairpin loop, interior loop, multiloop, stem within these RNA-sequences. To assess enrichment of significant edQTLs within RNA-secondary structures, for each structure, we randomly selected non-significant eQTLs in equal number of significant edQTLs in each structure category, which were matching minor allele frequency (MAF) and distance from edit sites with 50% standard error of both features for 1,000 times. We computed the enrichment p-value for each RNA structure as a number of observations in which the overlap of nonsignificant edQTLs with the RNA structure was higher than the overlap of significant edQTLs with the RNA structure divided by 1,000.

### Comparison of molecular QTLs

We extracted cell-type-specific primary eQTLs and sQTL from our previous study^26^, and cell-type-specific primary edQTLs and apaQTLs discovered in the current study for comparison. For distance from TSS/TTS, we calculated the distance between genetic variant and TSS/TTS of genes for which expression was tested in eQTLs, the distance between genetic variant and TSS/TTS of genes where alternative splicing tested was located in sQTLs, the distance between genetic variant and TSS/TTS of genes where edit site tested was located in edQTLs, and the distance between genetic variant and TSS/TTS of genes where APA site tested was located in apaQTLs considering the expression of the gene in either forward or reverse strand. For distance from splice sites, we computed the distance of genetic variants from either intron start and end sites for all potential alternative splicing events for a given gene, and used the shortest distance for comparison. To compare enrichment of primary QTLs within chromatin accessibility sites, we assessed the overlap of genetic variants within chromatin accessibility regions which were differentially accessible in progenitors/neuron^70^ for progenitor and neuron QTLs, respectively. To compare enrichment of primary QTLs within RBP binding sites, we utilized CLIPdb data, and assessed the overlap of genetic variants with this dataset^88^. Pairwise comparisons were performed via chi-square test.

### GWAS colocalization analysis

We applied LD-thresholded colocalization analysis to find edQTLs and apaQTLs colocalized with the traits for each cell type separately^26,125^. Summary statistics from GWAS for schizophrenia (SCZ)^3^, educational attainment (EA)^93^, major depression disorder (MDD)^126^, cortical thickness and surface area from UKBB^6^, and the ENIGMA project^5^, neuroticism^127^, IQ^4^, cognitive performance (CP)^93^, bipolar disorder (BP)^1^, attention-deficit/hyperactivity disorder (ADHD)^128^, Parkinson’s disease (PD)^129^ and Alzheimer’s disease (AD)^130^ were used. Index GWAS SNPs were defined as two LD-independent genome-wide significant GWAS signals (p-value < 5×10^-8^) with pairwise LD r^2^ < 0.2 calculated by using European population of 1000 Genomes (phase 3). For comparison of colocalizations with eQTLs and sQTLs^26^, we used LD r^2^

< 0.5 to detect index GWAS variants. To perform colocalization analysis, first, we detected two variants (one from index variants of GWAS and one from index variants of the QTL study) which had pairwise LD r^2^ greater than 0.8 based on either European population or our study). Then, we re-performed the edQTL/apaQTL analysis by conditioning on GWAS index variant, and if the association between edit rate/APA usage and the QTL index variant was no longer significant, we considered these two loci as co-localized.

