## Supplementary file for "Genetics of cell-type-specific post-transcriptional gene regulation during human neurogenesis"

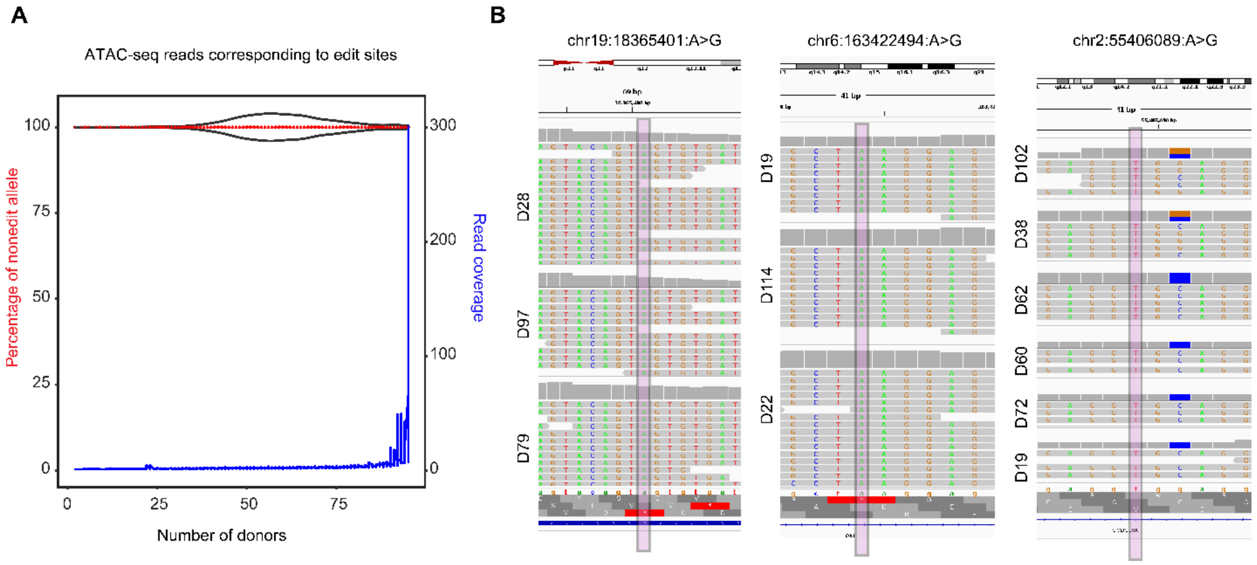


**Figure S1. Validation of RNA-editing sites discovered in RNA-seq data via ATAC-seq data.**

(A) The number of donors supporting that genomic DNA includes unedited alleles are shown on the x-axis. The percentage of unedited allele per edit sites is shown on the right y-axis (red) and the total number of ATAC-seq supporting unedited allele is given on the left y-axis (blue).

(B) Example ATAC-seq reads for RNA-editing sites chr19:18365401:A>G, chr6:163422494:A>G and chr2:55406089:A>G.

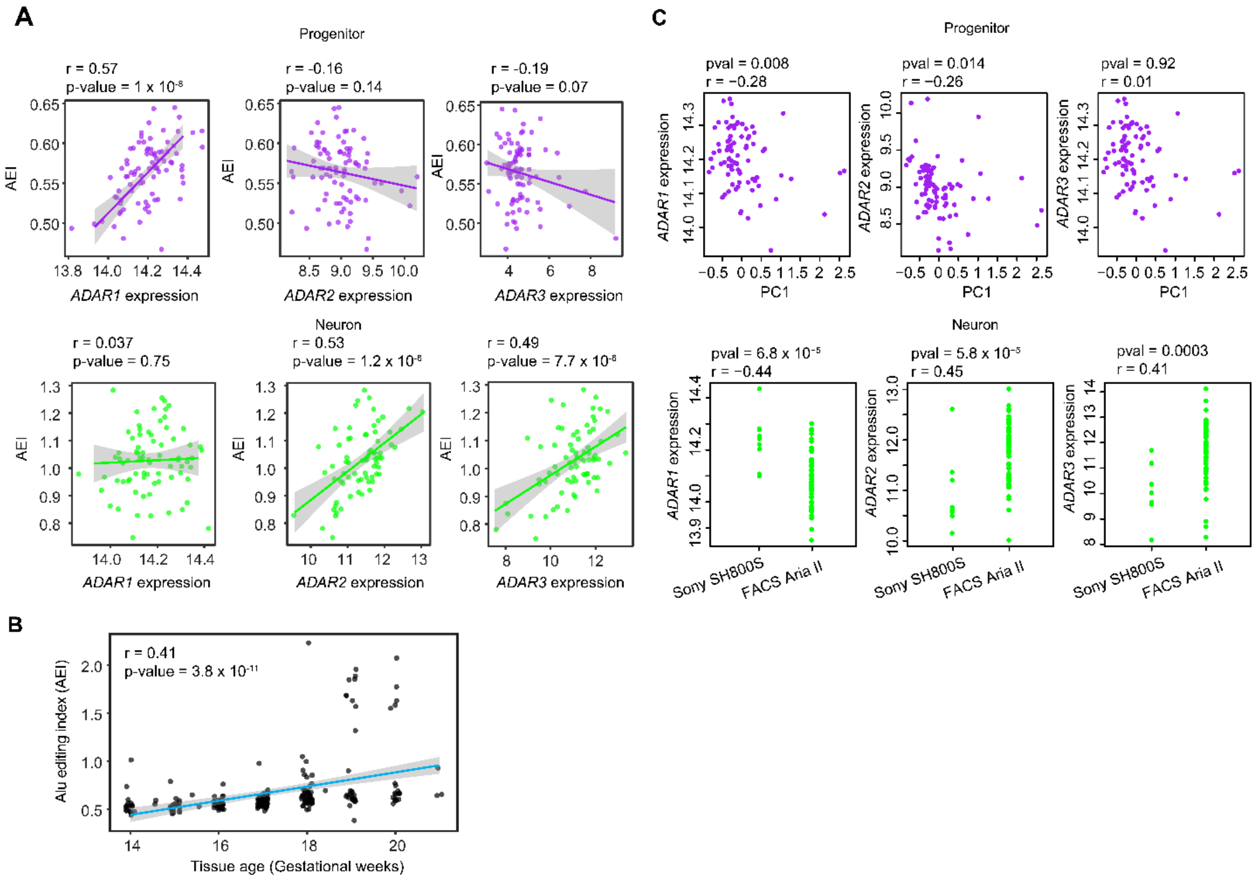


**Figure S2. ALU editing index is correlated with ADAR gene expression and developmental time.**

(A) Correlation of AEI values with VST normalized ADAR1, ADAR2 and ADAR3 expressions in progenitors (purple) and neurons (green). Correlation coefficients (r) and p-values are shown per relationship.

(B) Alu editing index across different gestational weeks in fetal bulk brain data.

(C) Correlation of technical confounders with *ADAR1-3* gene expressions per cell-type. Correlation efficient (r) and p-values are reported.

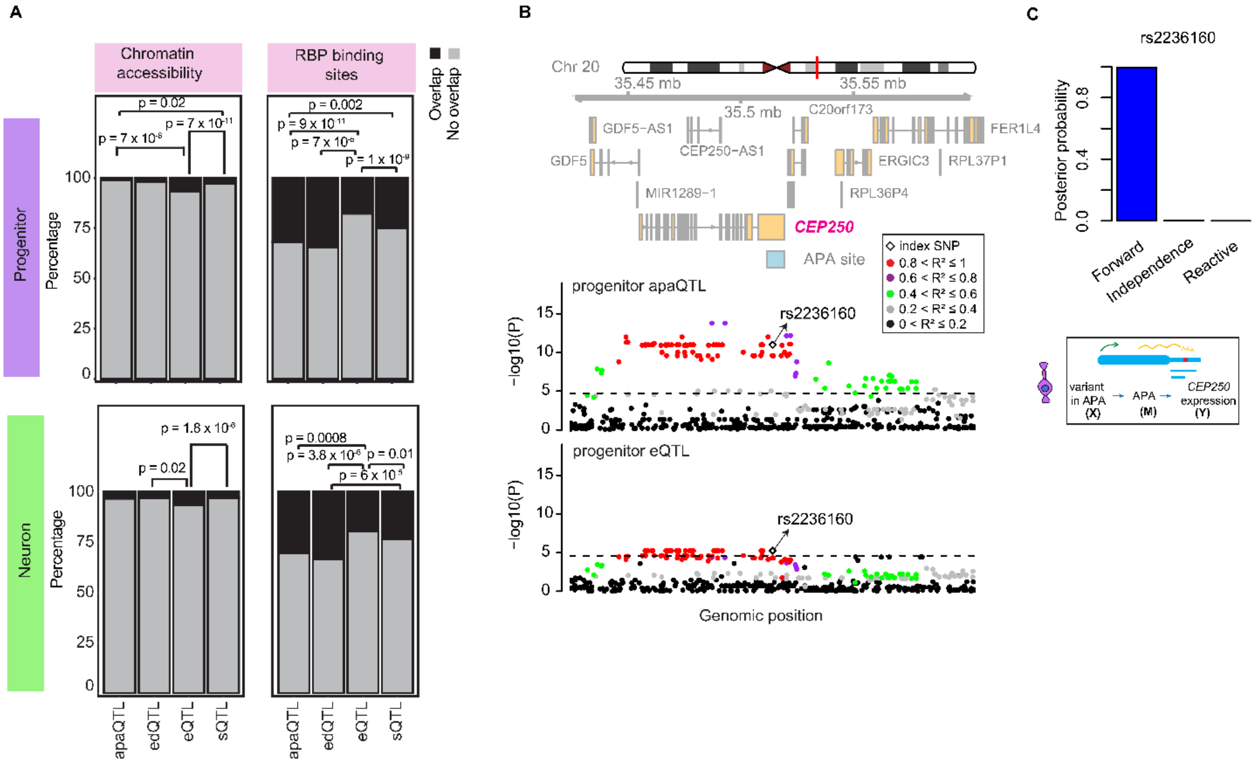


**Figure S3. Comparison of cell-type-specific molecular QTLs**

(A) Overlap of primary eQTLs, sQTLs, edQTLs and apaQTLs with chromatin accessibility regions are RBP binding sites per cell type. Chi-square test p-values (p) are reported for each pairwise comparison.

(B) APA site (chr20:35511626-35519280) within the *CEP250* gene mediated its expression. Genomic tracks for progenitor eQTLs and apaQTL are shown, and tracks are colored based on relative LD $r^2$ for the variant rs2236160, which is the nearest variant to APA site.

(C) Posterior probability of forward, independence and reactive models for mediation of *CEP250* gene expression via APA site. Lower cartoon illustrates the relationship between variant (X) within the APA site, mediator APA site (M) and *CEP250* gene.

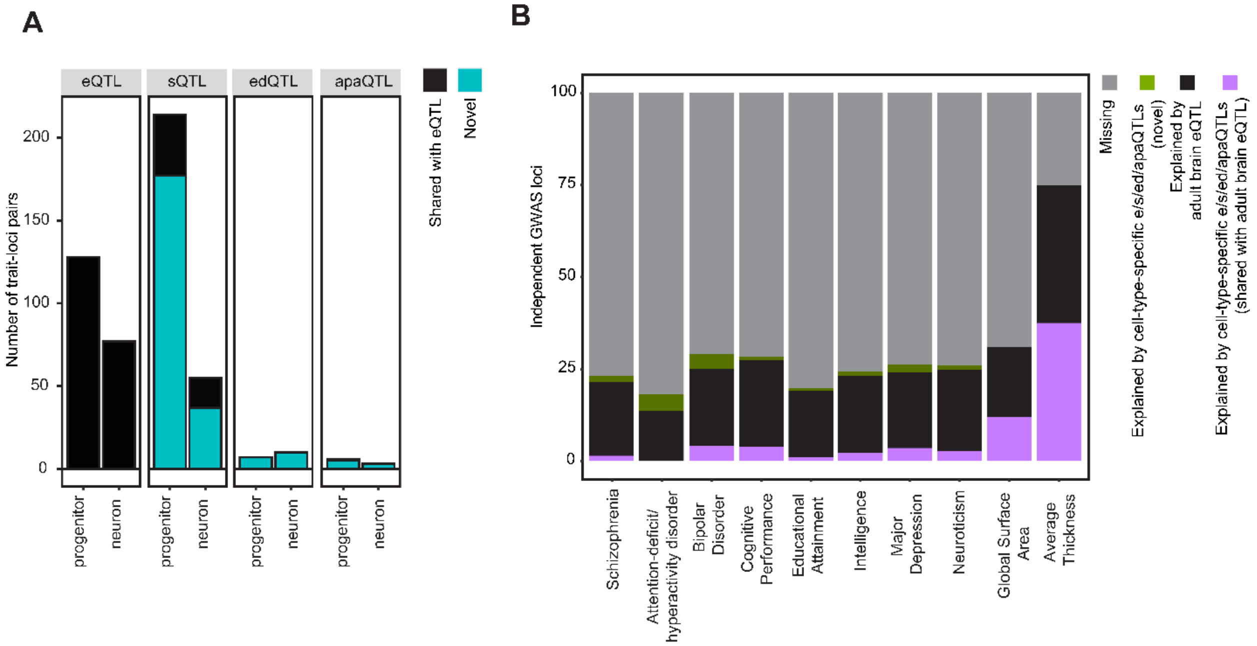


**Figure S4. Comparison of GWAS colocalizations across cell-type-specific molecular QTLs**

(A) Number of brain-relevant trait-loci pairs are found in eQTLs, sQTLs, edQTLs and apaQTLs, and shared pairs with eQTLs are indicated by black color, and pairs which were detected by using QTLs other than that eQTLs are indicated by blue color.

(B) Number of brain-relevant trait-loci pairs are found in cell-type-specific eQTLs, sQTLs, edQTLs and apaQTLs in addition to loci explained by adult brain eQTLs.

**Supplementary table legends**

**Table S1. RNA-editing sites discovered per cell type**

Edit: editing site (chromosome_genomic position_nonedit_editted base); strand: strand of the gene harboring edit site; Rep: overlap with repeats; shared_brainvar: overlap with BrainVar data; shared_gtex: overlap with GTEx Cortex; gene: gene harboring edit site; region: the genomic region including edit site; type: the data type exhibiting edit site.

**Table S2. Correlation of editing levels in neurons with TUJ1 + cells**

Edit: editing site (chromosome_genomic position_nonedit_editted base); r: correlation coefficient; pval: correlation pvalue; FDR: false discovery rate.

**Table S3. edQTLs in progenitor, neuron and fetal bulk data**

Snp: variant; edit: editing site (chromosome_genomic position_nonedit_editted base); beta: beta coefficient; pvalue: p-value; gene: gene harboring edit site; type: the data type exhibiting edit site; A1: effect allele.

**Table S4. APA sites discovered per cell type**

FilterID: alternative 3’UTR (chromosome_genomic position_3’UTR start site_3’UTR end site_strand); Gene: the gene harboring 3’UTR; Transcript: the isoform harboring 3’UTR; type: the data type exhibiting 3’UTR.

**Table S5. apaQTLs in progenitor, neuron and fetal bulk data**

Snp: variant; beta: beta coefficient; pvalue: p-value; utr: alternative 3’UTR (chromosome_genomic position_3’UTR start site_3’UTR end site_strand); gene: gene harboring edit site; type: the data type exhibiting edit site; A1: effect allele.

**Table S6. GWAS colocalization with cell-type-specific edQTLs and apaQTLs**

**First sheet:** Snp: variant; edit: editing site (chromosome_genomic position_nonedit_editted base); beta: beta coefficient; pvalue: p-value; chr: chromosome; cond.snp: GWAS variant; cond.beta: beta coefficient after conditioning; cond.pval: p-value after conditioning; trait: GWAS trait; type: the data type exhibiting edit site; GWAS_clump_r2: LD $r^2$ value used for colocalization analysis.

**Second sheet:** snp: variant; beta: beta coefficient; pval: p-value; poly: alternative 3'UTR (chromosome_genomic position_3'UTR start site_3'UTR end site_strand); GWASsnp: GWAS variant; Condbeta: beta coefficient after conditioning; Condpval: p-value after conditioning; r2: LD r2 between QTL and GWAS data; trait: GWAS trait; type: the data type exhibiting 3'UTR; GWAS_clump_r2: LD $r^2$ value used for colocalization analysis.
